# A predictive epilepsy index based on probabilistic classification of interictal spike waveforms

**DOI:** 10.1101/373571

**Authors:** Jesse A. Pfammatter, Rachel A. Bergstrom, Eli P. Wallace, Rama K. Maganti, Mathew V. Jones

## Abstract

Quantification of interictal spikes in EEG may provide insight on epilepsy disease burden, but manual quantification of spikes is time-consuming and subject to bias. We present a probability-based, automated method for the classification and quantification of interictal events, using EEG data from kainate- and saline-injected mice (C57BL/6J background) several weeks post-treatment. We first detected high-amplitude events, then projected event waveforms into Principal Components space and identified clusters of spike morphologies using a Gaussian Mixture Model. We calculated the odds-ratio of events from kainate-versus saline-treated mice within each cluster, converted these values to probability scores, P(kainate), and calculated an Hourly Epilepsy Index for each animal by summing the probabilities for events where the cluster P(kainate) > 0.5 and dividing the resultant sum by the record duration. This Index is predictive of whether an animal received an epileptogenic treatment (i.e., kainate), even if a seizure was never observed. We applied this method to an out-of-sample dataset to assess epileptiform spike morphologies in five kainate mice monitored for ~1 month. The magnitude of the Index increased over time in a subset of animals and revealed changes in the prevalence of epileptiform (P(kainate) > 0.5) spike morphologies. Importantly, in both data sets, animals that had electrographic seizures also had a high Index. This analysis is fast, unbiased, and provides information regarding the salience of spike morphologies for disease progression. Future refinement will allow a better understanding of the definition of interictal spikes in quantitative and unambiguous terms.

## Introduction

The electroencephalogram (EEG) is an essential tool for monitoring seizures, diagnosing epilepsy and understanding epileptogenesis. Most EEG analysis, including the scoring of seizures and interictal epileptiform events, is carried out manually by trained scorers. Manual scoring is labor-intensive and prone to low interobserver agreement (1) which often precludes fine-scale and quantitative analysis of EEG, especially for chronic (>24 hours) recordings commonly employed in clinical and research settings (2,3).

Adding to the challenge of EEG analysis and diagnosis in epilepsy is a relatively low seizure frequency, often much less than one/day, making capturing confirmatory seizure events on EEG rare (4). Thus, the identification and analysis of subclinical, interictal events such as spikes may be important for diagnosis, identification of epileptic foci and understanding the progression of epilepsy (3-10). But the definition of spike-like events for subsequent manual scoring remains an area of some disagreement among clinicians (2) and researchers (1,11). Together with the laborious nature of manual EEG analysis, this means that the frequency of interictal spiking is not typically quantified for diagnostic purposes in the clinic, though there are special cases, and is underutilized in epilepsy research.

There are many automated approaches for detecting interictal events, including template-matching (12), non-linear energy operator (NEO) detection (13,14) wavelet filtering and line length (15), and neural networks (16) among others (17). These algorithms rely in part upon a spike definition that is based on expert visual analysis and are rooted in some version of the definition by Chatrian et al 1974, where a spike is a transient that is clearly distinguishable from background activity, with a duration from 20 to 70 ms (18-20). Still, a universal ground truth definition of an interictal spike is lacking as demonstrated in many cases where interobserver agreement may be relatively low (1,21). Further careful analysis of spike and spike-like event morphology is needed to refine this definition and to improve the utility of spike analysis in epilepsy.

Animal studies also demonstrate that spiking of some type can be present in both normal and epileptic brains of rodents (2,4,22,23), making it important to identify aspects of spike-like morphologies that are specifically associated with epilepsy. But differentiation of the many spike types and shapes by visual or algorithmic analysis is a difficult prospect. Rigorous differentiation and quantification of spikes in epileptic versus normal animals could elucidate patterns of activity that shed light on spike-related mechanisms in epilepsy and identification of epileptic foci while also contributing to an unbiased definition of true interictal spiking activity across models of epilepsy. Given the links between interictal spiking and disease progression in epilepsy (24,25), quantitative understanding and differentiation between normal and epileptiform spike-like activity is essential for future progress in epilepsy research; studies in animal models are essential to this progress.

Classification of spikes based on waveform morphology is a promising approach for differentiating events that are normal versus those that are relevant to epilepsy. Indeed, spike sorting by waveform clustering has been used to tune spike-detection and quantification algorithms (26), to choose spike templates (27) and to identify or predict epileptic foci (28). A very well-studied use of waveform clustering is to distinguish individual extracellularly recorded action potential waveforms in order to identify neurons in single-unit recordings (29). Several clustering approaches have been used across these applications, including K-means clustering and affinity propagation (27), graph theoretical algorithms (28), and Gaussian Mixture Models (GMM) (29).

Here we present a novel waveform-based analysis to compare spike-like events recorded from mice treated with an epileptogenic insult (i.e., kainate, a model of temporal lobe epilepsy (30)) with spike-like events recorded from control mice treated with saline. We identified spike-like events in both groups, then projected these waveforms into a low-dimensional (3D) space using Principal Components Analysis (PCA), and clustered these waveforms using a GMM. This allowed us to estimate the probability distributions that characterize normal versus epileptiform spike-like waveforms and to compute the probability that specific spike waveforms reflect epileptiform activity. By assigning a probability score to each event type, we provide a novel measure (the Hourly Epilepsy Index) for estimating the probability that an animal is epileptic, even in the absence of observing electrographic or behavioral seizures. The ability to assess epilepsy with a high-resolution metric in every animal is extremely important in animal models with low seizure burden such as repeated low-dose kainite in C57Bl6 mice (used in this study) and could result in the more efficient use of animals and EEG recording time in future studies. This novel analysis thus provides rapid and unbiased insight into differences among spike types in normal and epileptic animals that can be used for early diagnosis of risk and serve as a biomarker of disease progression.

## Materials and methods

### Animals

All use of animals in this manuscript conformed to the *Guide for the Care and Use of Laboratory Animals* (31) and was approved by the University of Wisconsin-Madison Institutional Animal Care and Use Committee (IACUC, protocol M005629).

Animals (male C57BL/6J background, ~5 weeks old) were received from Harlan (Madison, WI) and housed in clear acrylic cages ~7 L cages with filtered airflow and corn cob bedding with paper nesting material. Animals were housed in groups of five or less with rodent chow and water available *ad libitum.* After ~2 weeks of acclimation, animals underwent repeated low-dose kainate (KA) or saline (SA) injections (~7 weeks old) and EEG implantation and recording (~18-20 weeks). All animals were checked daily for food/water/health by laboratory personnel and/or the University of Wisconsin-Madison Biomedical Research Model Services. Staff veterinaries were consulted in the event of illness (post-injection) or severe injury (as a result of fighting post-injection) and their recommendations for animal care were followed.

The data used in this study represents a portion of data collected as a result of an additional study (IACUC approved) investigating pneumonic discrimination during behavioral trials and computational pattern separation *in slice* (*In Preparation*).

### Epilepsy induction

During the injection process, when not handled, mice were individually housed in enclosed ~150 cm^3^ acrylic cubicles with opaque sides and clear front portals with holes to allow air exchange, and equipped with corn cob bedding and rodent chow. Animals were then randomly assigned to SA or KA treatment, ear punched or tagged for identification and weighed. Mice received a series of intraperitoneal (IP) injections based on the following schedule (herein referred to as “repeated low-dose kainate”).

Mice in the KA group first received a 10 mg/kg (5.5 mM) dose of kainate (Tocris Bioscience, UK) delivered in 1x phosphate-buffered saline (PBS) prepared from PBS tablets (Dot Scientific, Michigan) dissolved in deionized distilled water (DDW) and filter sterilized (0.22 µm, Corning, MI). Mice continued to receive injections at 2.5-5 mg/kg every 20 minutes until status epilepticus (SE) occurred (4-9 injections) (32,33). Mice in the SA group received the same schedule of sterile PBS injections. Mice were considered to be in SE when displaying behavioral seizures of level 4-5 on the Racine Scale (34) for a minimum of 30 minutes. Mice usually experienced SE for 90-180 minutes and were allowed to recover spontaneously. Mice were treated in cohorts of ~8 at a time, with a ratio of ~2:1, KA:SA animals. The first cohort of mice received alternating 5 mg/kg and 2.5 mg/kg injections after the initial 10 mg /kg dose, but this schedule took longer than only 5 mg/kg injections without any noticeable differences in eventual efficacy or survival, so subsequent cohorts received 5 mg/kg injections. Across all cohorts, mortality was <10%.

After the injection schedule, animals were given fresh apple slices, monitored until SE ceased (assessed by normal posturing), and returned to group housing. They were monitored, weighed, and given 0.4 mL of sterile 1x PBS via IP injection on a daily basis as necessary until their weight returned to pre-injection level. No animal injected with SA ever experienced status epilepticus, lost weight, or required recovery injections of 1x PBS. Mice remained in the vivarium for 7-9 weeks after injections, allowing time for the development of epilepsy in KA mice. During this time some KA mice became aggressive and were removed from group housing and caged individually.

### EEG surgery and recording

Animals were implanted with EEG headcaps while under 1-2% vaporized isoflurane anesthesia (see (35) for detailed methods) at ~18-20 weeks old (~11-13 weeks post injection), allowed to recover for 3-4 days, and then recorded continuously at 256, 512, or 1024 Hz sampling rate for three days. All data not collected at 512 Hz were either resampled (1024 Hz) or linearly interpolated (256 Hz) to 512 Hz for analysis. Data were then notch filtered at 60 Hz using a Chebyshev Type II digital filter and high-pass (>0.5 Hz) filtered with a Chebyshev Type I digital filter. All analyses presented in this manuscript used a right-frontal EEG channel.

### High-amplitude event detection

To assess the severity of epilepsy in animals injected with kainate, we developed a probability-based approach for the detection of epilepsy-related events (e.g. interictal abnormal events like EEG spikes, seizures) and the calculation of an epilepsy severity index.

Filtered EEG signals were normalized to using a variation of z-score normalization that we term Gaussian normalization. First, we fit a Gaussian distribution to the all-points histogram of each 24-hour record using the Nelder-Mead Simplex (36) method as implemented by the fminsearch function in Matlab. We then normalized the 24-hour record by subtracting the mean from each point and dividing by the standard deviation of the model fit. The resultant normalization is different from the standard z-score normalization in that the mean and standard deviation components are calculated from a model estimating the Gaussian components of the EEG rather than empirically calculating the mean and standard deviation. We employed this normalization because it adjusts the data based on the normal component of the signal and is less influenced by very large and brief artifacts.

Using normalized EEG we detected high-amplitude, spike-like events using a two-threshold strategy (37). The final thresholds used in this analysis were optimized to yield the largest predictive effect size (Fig 1D). The start of an event was triggered when the EEG signal rose 5 standard deviations above the mean of the Gaussian fit. An event ended when the signal passed 1 standard deviation below the mean. We then combined any events occurring within 0.2 seconds of one another (end to start) into a single event (19). This resulted in a list of events of varying duration (98% shorter than 1s, with some representing rapid spike bursts) from each animal/day EEG record. We then took two seconds of signal centered on the start of each event and collated events across 72 days of recording (3-4 days from each of 15 KA and 7 SA animals). This resulted in a matrix with ~33000 event waveforms (1024 points each) across all treatments, recording days, and animals. We chose optimal high (event start) and low (event end) threshold values after a grid search to investigate high values between 3 and 8, and low values between −3 and 2 and then performing a t-test to compare the number of events per hour identified in SA vs. KA animals (Fig 1D).

### Epileptiform event confirmation

We compared the output of the two-threshold detection method with output from another method that has been highly cited for seizure and interictal spike detection, and that also performs similarly to manual human scoring (Bergstrom et al. 2013). Spikes, seizures, and other abnormal epileptiform activity were detected in normalized EEG according to Bergstrom et al. (2013). The baseline was determined using the whole signal. The event threshold (SF value) was set at 4.0, where spikes were defined using a modified amplitude of 15, and seizures were defined as events greater than one second in duration. To compare the two-threshold method with that of Bergstrom et al. (2013), all records were represented as logical vectors where 1 indicates the presence of an event during each second of recording and 0 indicates the absence of an event. Confusion matrices (38) between these vectors were used to compute true-positive and false-negative rates, by provisionally assuming that Bergstrom et al. (2013) detected “true” events.

### PCA, clustering, and P(KA)

We performed Principal Components Analysis (PCA) (39,40) on the matrix (~33000 x 1024) of waveforms detected by the two-threshold method and projected each original waveform into the space spanned by the first three Principal Components (PCs, Fig 3). In such 3D projections, each point represents a detected waveform and the distance between any two points represents the difference between their original waveform shapes. Three dimensions were chosen for ease of display, but higher or lower dimensional projections could be used throughout this analysis in principle.

Waveforms that are closely intertwined with epilepsy should be a) more common in KA than in SA and b) similar to other waveforms in epileptic (KA) mice and different from those of SA mice. Therefore, the densities of PCA-projected points for KA and SA events are expected to overlap somewhat but, by examining quantitatively their overlap in specific regions of PCA space, we can compute the likelihood of a waveform in any region having come from a KA vs SA animal.

The resulting ensemble of points was fit with a Gaussian Mixture Model (GMM) with expectation maximization using the function fitgmdist() (41) with nine components and then clustered based on the GMM coefficients. The GMM defines a probability for every point in the space, and in this case reflects the prior probability distribution that any waveform would be detected because both KA and SA were included in the GMM fit. We then calculated the conditional probability, P(KA) that events in each cluster are related to an epileptogenic treatment by computing the ratio of KA (epileptogenic) to SA (control) events in each cluster (Odds ratio) and converting it to a probability, P, using 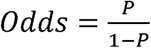.

### Hourly Epilepsy Index

Given that there is some disagreement between manual scorers, we are attempting to provide a statistical and unbiased measure of epileptiform event frequencies. We first defined epileptiform event morphologies as events with a P(KA) > 0.50, because they are more likely to be found in KA than SA animals and are therefore more relevant to epilepsy. The range of P(KA) values was rescaled from 0.5 - 1.0 to 0 - 1, where *scaled P*(*KA*) = (*P*(*KA*) - 0.5) × 2 to reflect the relative contribution of each event morphology to the KA-specific set. Our method is probabilistic, so it can only be interpreted as an “index” rather than an actual count of interictal events. Therefore, we define the *Hourly Epilepsy Index* = ∑(*freq. events in each cluster with P(KA)* > 0.5 × *associated scaled P(KA)*).

### Parameter optimization

One tunable parameter in this analysis is the choice of how many clusters to use in modeling the data (i.e., how fine-grained should our analysis of waveform shapes be?). To investigate this parameter, we built a set of models varying the total number of GMM clusters. Additionally, because creation of a GMM in our implementation is a stochastic process, we rebuilt each model at 25 unique random seeds (seeds = 1 through 25). In total, we generated 600 Gaussian mixture models and related cluster profiles, P(KA)s, and Hourly Epilepsy Indices. From each of these model sets we calculated: 1) the Negative Loglikelihood of the data fit to the GMM, 2) The average belongingness of each event to its assigned cluster (Cluster Belongingness, defined below), 3) the effect size between SA and KA treated animals and 4) *P* value comparing the Hourly Epilepsy Index between SA and KA treated animals. Cluster Belongingness was calculated by taking the sum of squares of the set of posterior probabilities, calculated with the posterior() function in Matlab, for each event and then averaged across all events. For each event, these posterior probabilities (as many probabilities as there are clusters) provide the likelihood of belonging to each cluster in the model and these probabilities sum to one. For example, in the case of a 2 cluster model, data point A may belong equally to each cluster *P* = (0.5, 0.5) where its Cluster Belongingness equals 0.5. In contrast, data point A may belong much more strongly to one cluster over another *P* = (0.9, 0.1) with a cluster belongingness of 0.82. Thus, a clustering model with poor fit will have a low value for Cluster Belongingness.

### Computer programming

All analyses, data processing, and event classification was done in Matlab (Mathworks, Natick, MA) using home-written scripts. The final version of this algorithm was executed with Matlab 2017b.

### Statistical analysis

The usage of statistical tests and procedures used to estimate model fit are indicated throughout the methods section of this manuscript. All tests and statistical applications were chosen after careful consideration of data distribution (e.g. normality) and independence. All statistical tests were two-tailed. The researchers were not blinded or randomized to the treatments given to animals, but rather treatments (KA and SA) known to cause the development of epilepsy (32,33) were used to calculate probabilistic information that went into creating the Hourly Epilepsy Index. Indeed, the design of this approach was to optimize for differences between the treatments. However, we feel that this approach is warranted and importantly, the clustering of the events morphologies was unsupervised and thus ‘blinded’ to the treatment from which each event originated.

### Data Availability

Code for the algorithm used in this manuscript has been made publicly available at the GitHub repository github.com/jessePfammatter/detectIISs. Code for the Bergstrom *et al.* algorithm can be found at github.com/bergstromr/IISdetection. Data are available in the BioStudies database (http://www.ebi.ac.uk/biostudies) under accession number S-BSST180.

## Results

### EEG recording and event detection

We collected three days of continuous, 24-hour EEG for 15 KA and 7 SA animals and normalized the records as described above. The two-threshold event detection algorithm identified 25681 and 5764 events in KA- and SA-treated animals, respectively (Fig 1; mean ± SD event duration for KA animals = 55 ± 148 ms; for SA animals = 169 ± 380 ms). The average number of events per hour detected in KA (23.3 ± 27.4) and SA (11.5 ± 11.0) animals using this detection method was not significantly different (Fig 1C, *P* = 0.166, *t* = 1.44, df = 19.83, two-sample T-test with unequal variances). The p-value for comparison of KA and SA animals was minimized by the use of the high threshold of +5 and low threshold of −1, though combinations of high thresholds of four or five and low thresholds of one to -2 showed low p-values relative to all other combinations (Fig 1D).

**Fig 1.**
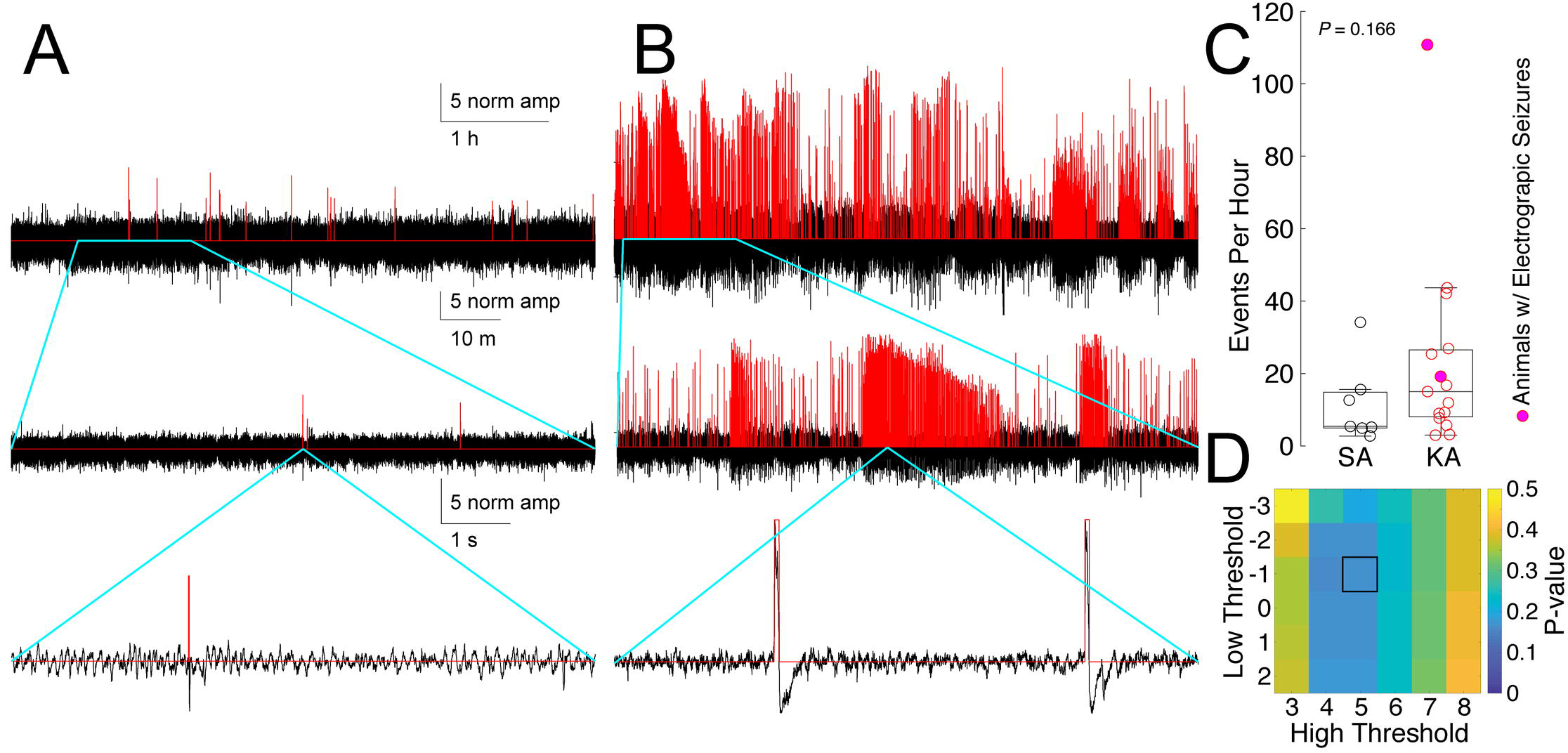
Event detection by two-threshold approach. Sample traces of events identified (red) using the two-threshold method (high threshold = 5 SD above Gaussian mean, low threshold = 1 SD below mean) in (A) SA- and (B) KA-injected animals. Each row in A and B shows an expanded view (indicated by the blue lines) of the signal directly above. (C) Average number of events per hour detected in SA (black) versus KA (red) animals (*P* = 0.166, *t* = 1.44, df = 19.83, two-sample T-test with unequal variances). Circles represent the average number of events per hour for each animal over 3 days of recording. Closed circles represent animals that were recorded having an electrographic seizure. (D) *P*-values comparing the average number of events per hour in SA and KA animals across a set of high and low thresholds; the p-value is minimized at 5 and -1. The center mark of the box and whisker plot represents the median, while the lower and upper bounds of the box represent the 25^th^ and 75^th^ quartiles, respectively; the whiskers extend to the most extreme data points or 1.5 × the interquartile range.

### High-amplitude events are epileptiform

To confirm that the events identified by the two-threshold method include epileptiform events, the normalized EEG signal was analyzed using an event detection and sorting algorithm that is validated with human spike scoring (15). A comparison between events detected by the two-threshold method and spikes from Bergstrom *et al.* resulted in a true positive rate (TPR) of 0.781 and a false positive rate (FPR) of 0.005 (Fig 2A, 2C). Thus, our initial detection of spikes agrees reasonably well with a previous method that itself agrees well with human scoring. In a comparison of the two-threshold detection method to all epileptiform events (spikes, seizures, and other low-amplitude, short duration abnormal) from Bergstrom *et al.* (2013), the agreement between the algorithms drops precipitously (Fig 2B, 2D, TPR and FPR for all events are 0.038 and 0.001, respectively), suggesting that the two-threshold method is selective for spike-type events and does not detect low-amplitude abnormal or some seizure-like events. We illustrate specific examples of agreement (Fig 2E) and disagreement (Fig 2F) between the two detection methods, highlighting that the two-threshold detection method selects for high-amplitude spike- and seizure-like activity.

**Fig 2.**
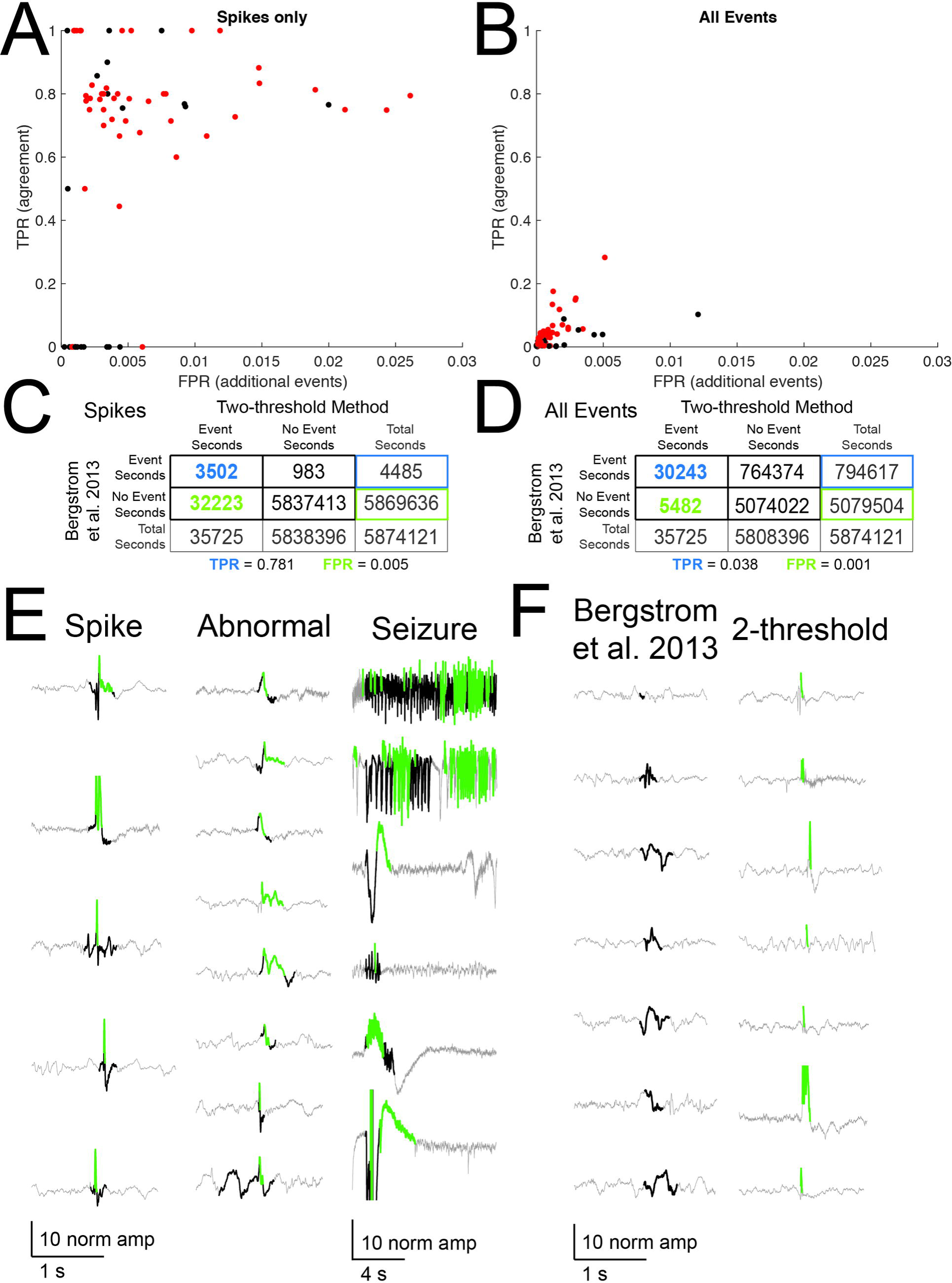
Interictal events correspond to epileptiform activity. We compared events from the two-threshold crossing method to events detected by a validated event detection and classification algorithm (15). Second-by-second, animal-specific (SA = black, KA = red) event classification agreement (True positive rate, TPR) and the rate of detection of additional events not identified in the validated method (Additional events, false positive rate, FPR) for spike-type events only (A) and all event types (seizures, spikes, and other low-amplitude abnormal and epileptiform events, (B), with corresponding confusion matrices (spikes only C, all events D). True positives are events detected by both the validated algorithm and the two-threshold crossing method. (E) Sample true positive events for each event type, where gray is the original EEG trace, green is two-threshold detection output, and black is Bergstrom *et al*., 2013 detection output. (F) Sample events for each detection method that were not detected by the other method.

### Measures predictive of an epileptogenic insult

Events from KA- and SA-treated animals were projected into Principal Components space (Fig 3A, first three PCs shown). We used the first three PC (76.7% of the overall variance) and clustered the data into nine groups using a GMM (Fig 3B). The probability, P(KA), that a cluster morphology is relevant to epilepsy is equal to the probability of that morphology being found in KA but not SA records and was calculated as the odds ratio of the number of KA to SA events for each cluster. The P(KA) for four of nine clusters is above 0.5 (Fig.3C, range 0.73 – 0.99), indicating that event morphologies in these clusters may be specific to the KA treatment and thus related to epilepsy. On average, KA animals exhibit a significantly higher average Hourly Epilepsy Index than do SA animals (11.6 ± 14.6 for KA, 3.0 ± 2.0 for SA; *P* = 0.039, *t* = 2.26, df = 15.04, two-sample t-test with unequal variances). Two of 15 KA animals displayed electrographic seizure activity in at least one recording (Fig.3D, magenta filled circles); the average Hourly Epilepsy Index for these animals falls above the 75th percentile (top line of box plot) of the SA hourly index. We found that there was a broad spread of Hourly Epilepsy Indices across the 17 KA mice, where eight KA mice had Hourly Epilepsy Indices similar to SA mice and 9 KA mice had Hourly Epilepsy Indices above the 95% confidence interval of the SA Index (95% confidence interval for SA = 1.14 - 4.47). This interesting result is in line with published reports of varying sensitivity to systemic KA injection in epileptogenesis (32,33).

**Fig 3.**
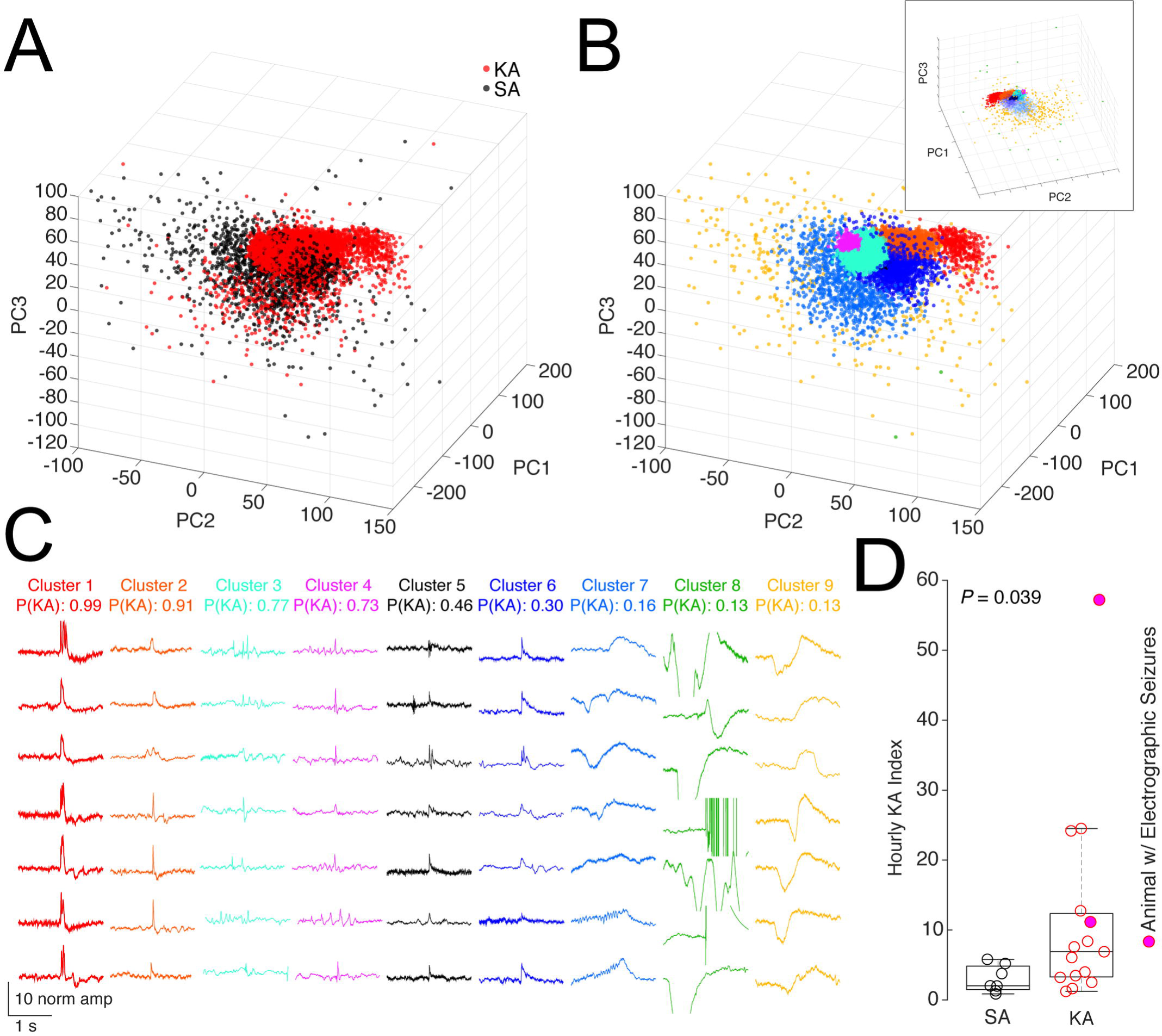
Event sorting by clustering and Hourly Epilepsy Index.

(A) Events detected with the two-threshold crossing method, from all animals and records, and projected into Principal Components space (first three PCs shown, red = KA, black = SA). (B) Events in A clustered into 9 groups using a GMM with Expectation Maximization. The upper inset shows a rotated and expanded view of the same plot, allowing a view of cluster 5 (embedded within cluster 6, which is set to 10% transparency for visualization). (C) P(KA) for each cluster identified in B calculated from the relative proportion of KA to SA events, presented in descending order with examples of events. (D) The average Hourly Epilepsy Index (15 KA, 7 SA animals) is much higher for animals treated with KA (*P* = 0.039, *t* = 2.26, df = 15.04, two-sample T-test with unequal variances). The center mark of the box and whisker plot represents the median, while the lower and upper bounds of the box represent the 25^th^ and 75^th^ quartiles, respectively; the whiskers extend to the most extreme data points or 1.5x the interquartile range.

### Optimization of the clustering algorithm

To determine the optimum number of clusters to differentiate between KA- and SA-specific spike morphologies, we calculated 600 repetitions of the Gaussian Mixture Model used to cluster events at each unique value of 2 to 25 clusters and random seeds set from 1 to 25. The Negative Loglikelihood (Fig 4A) approaches an asymptote around eight to ten clusters, suggesting the reasonable model fit in this range. Cluster Belongingness (Fig 4B) decreases across the simulation range with no clear inflection point (i.e. elbow) in the data to indicate diminishing returns of increasing cluster number. Effect size and p-value for the Hourly Epilepsy Index (Fig.4C and D) models indicate that the effect size and the p-value reach an asymptote in a range similar to that indicated by the Negative Loglikelihood results. Here, we have a fuzzy problem in terms of cluster selection. We selected nine clusters for our purposes and discuss cluster selection in depth in the discussion section.

**Fig 4.**
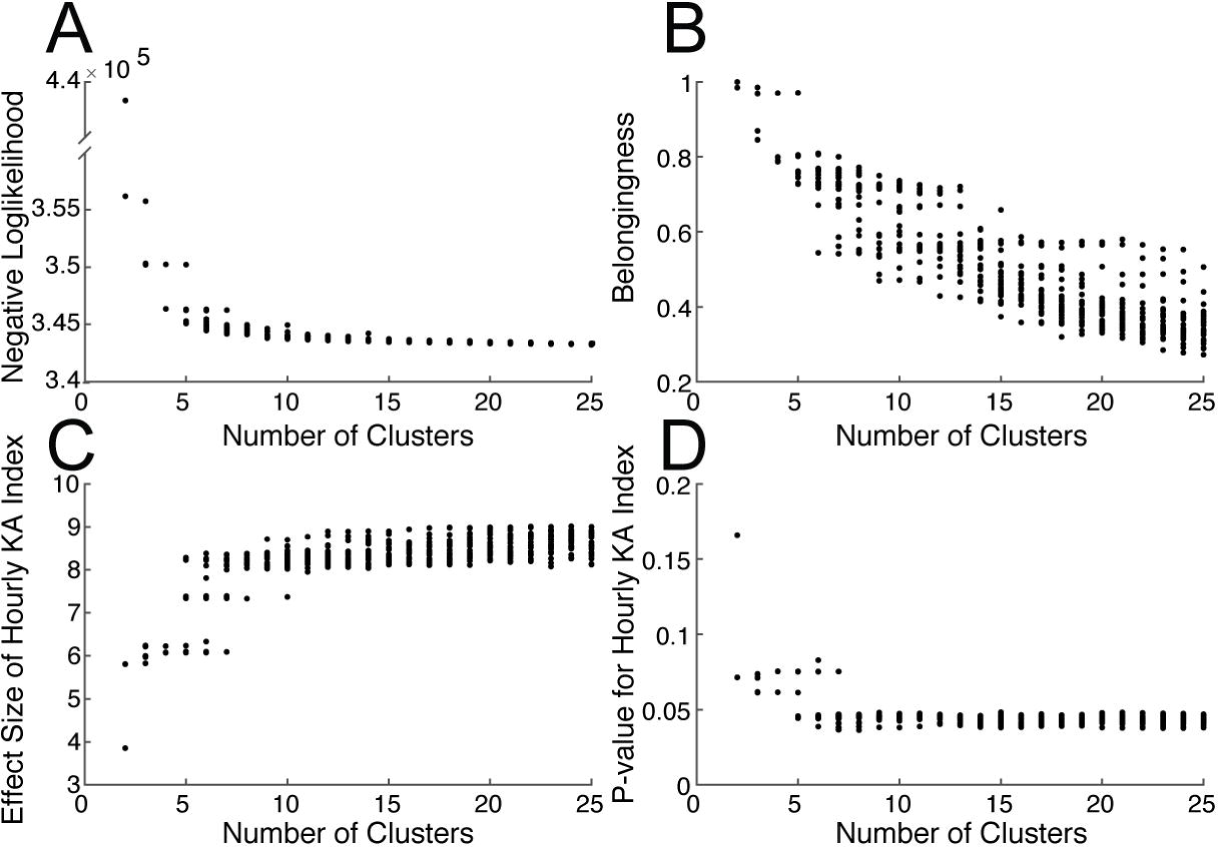
Selecting an appropriate number of clusters. We repeatedly generated a set of models (n = 25, random seeds set from 1:25) specifying a range of 2 to 25 clusters (totaling 600 models) and calculated the (A) Negative Loglikelihood, (B) Cluster Belongingness, the (C) Effect size and (D) *P* value of the Hourly KA Index comparing KA to SA treated animals (calculated using only events where P(KA) > 0.5). These values offer guidelines for model selection via ‘elbows’ in the data which are indicative of diminishing returns in model fit. It does not appear that there is a clear best value for clustering these data into discrete groups. Therefore, the cluster number must be selected and evaluated based on the goal of the cluster analysis.

### Cluster morphologies match published characteristics of interictal spikes

We performed a preliminary waveform shape analysis by plotting individual event duration vs amplitude for each cluster. In clusters where P(KA) > 0.5 (clusters one through four, Fig 5A) many events from KA-treated animals have a higher amplitude than do events from the same cluster found in SA-treated animals. The majority (92%) of events in these clusters for both treatment groups are shorter than 200 ms, consistent with published spike definitions. This trend is not as distinct in clusters five through nine, where P(KA) < 0.5. Further inspection of events in clusters one through four reveals that the morphologies of these event types are more classically “spikey” (short-duration events with sharp waveforms) than are events found in the remaining clusters. Finally, the majority of the events identified in KA animals (88%) were found in clusters one through four, whereas events in SA animals were equally distributed among the clusters with P(KA) above (49%) and below 0.5. The average number of events per hour for each cluster in SA and KA shows separation between event frequency in clusters one through four but not clusters five through nine (Fig 5B, p-values reported on each graph).

**Fig 5.**
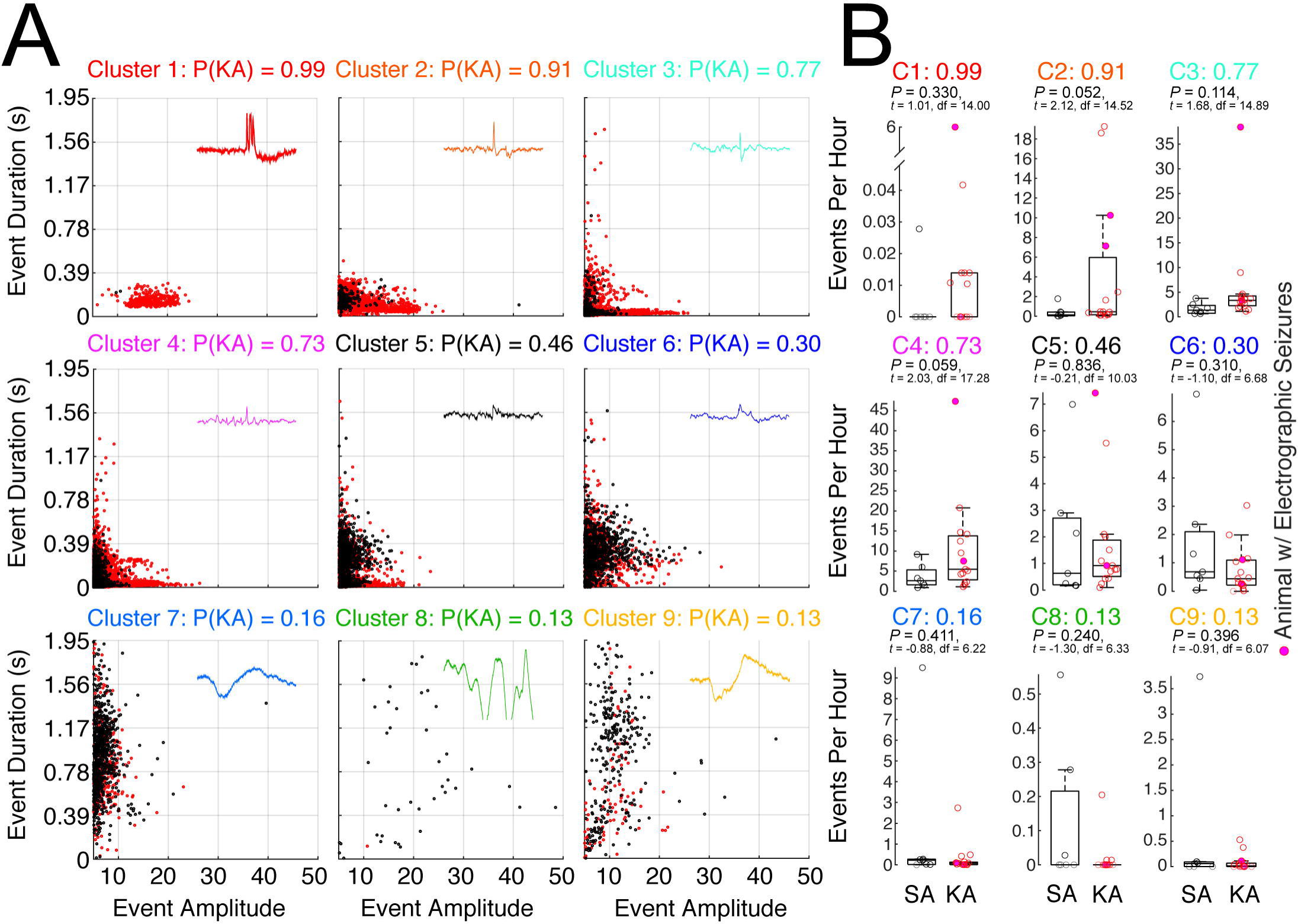
Characterization of individual clusters and contribution to total event count. (A) Graphical distribution of each cluster’s events in EEG amplitude (x-axis) by event duration (y-axis, time (s)) for events belonging to both KA (red) and SA (black). Example events from each cluster type are displayed in the upper right corner of each plot. For improved visualization of the majority of data, we have set the axis to exclude a small amount of the most extreme data points (most notably from cluster 7). (B) The average number of events per hour within each cluster for SA (black) and KA (red) animals. P-values indicate the results of t-tests between KA and SA for each cluster of waveform shapes.

### Analysis of out-of-sample data supports the methodology

We applied the clustering approach to events detected in a novel dataset of longitudinal recordings from 5 KA-treated animals (Fig.6A - D, recorded on days 30 – 34 and 44 – 51 post-KA). The longitudinal data were projected into the same PC space as the SA and KA dataset from Fig.3A and clustered according to the same parameters as Fig.3 (Fig 6A, B). All but one animal (Animal E) experienced similar, but not identical, increases over time in total event rate (average for all events per hour, Fig.6C) and Hourly Epilepsy Index (clusters one through four, Fig 6D). Electrographic seizures were only detected in one animal (Animal B) at days 44 and 45; this animal displayed a trend toward increasing event rate and Hourly Epilepsy Index (Fig.6C-E, magenta-filled circles).

**Fig 6.**
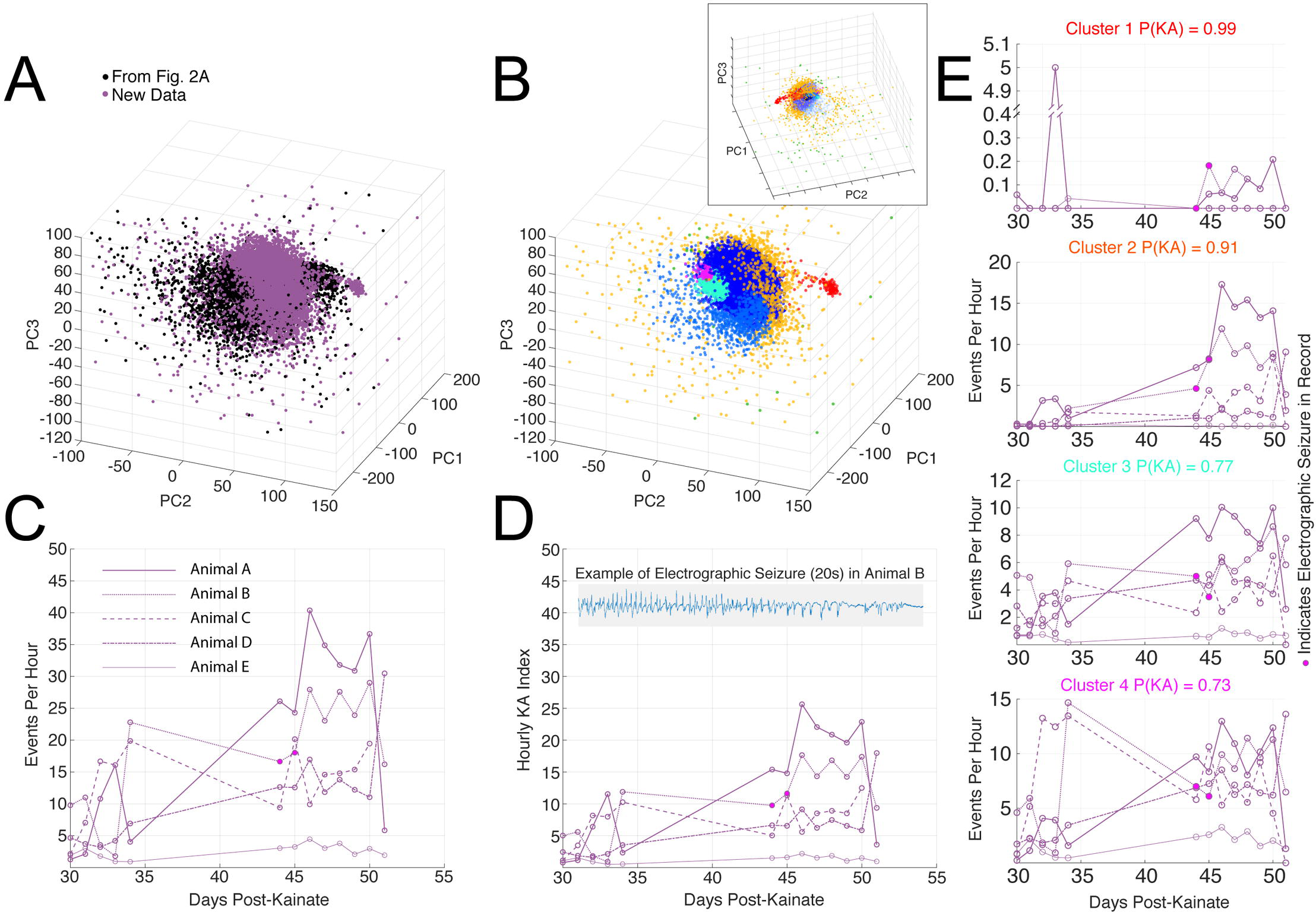
Detection and categorization of interictal events in longitudinal data. We applied our detection and categorization algorithm to longitudinal EEG records (recorded days 30 – 34 and 44 – 51 post-KA) from 5 KA mice. (A) Events identified from these 5 animals (purple circles) projected into PCs with data from Figure 3 (black circles, SA and KA data). (B) Clustering of these new data points using the same GMM coefficients and clustering parameters used in Figure 3 (cluster colors correspond to those throughout manuscript). The upper inset shows a rotated and expanded view of the same plot, allowing a view of cluster 5 (embedded within cluster 6, which is set to 10% transparency for visualization). (C) Events per hour as detected by the two-threshold method in each of the 5 KA animals over time. (D) Hourly KA Index for longitudinal data set. E) Average events per hour as detected by the two-threshold method within each cluster group. In panels C-E, records with electrographic seizures are shown as filled in circles. An example seizure is shown in the inset of panel D.

### Discussion

Interictal spikes are a hallmark of the epileptic brain (2). Yet the definition of an interictal spike remains somewhat nebulous, potentially biasing spike-detection strategies and algorithms. Previous analyses of spikes in kainate and other models of epilepsy have relied upon scoring of spikes and other interictal events to differentiate between KA-treated and control animals, where more spikes are observed in epileptic than in control animals (4,15). In these studies, control animals also exhibit spike-type events in the EEG; however, given that control (and many KA animals) animals do not proceed to have seizures, it is probable that some spikes arise by a non-epileptic mechanism, including spike-like events in normal brain (e.g., sharp wave ripples (2), electrographic noise, artifact, or damage associated with electrode implantation. That non-epileptiform spikes may also be present in both normal and kainate-treated animals further complicates the quantification of interictal spikes and thus the interpretation of nonconvulsive EEG events in epilepsy and epileptogenesis.

Here we present a novel automated algorithm for probability-based differentiation of epileptiform versus non-epileptiform spike and interictal EEG event morphologies in the kainate model of TLE. Our algorithm detects interictal events and weights them in an Hourly Epilepsy Index based on the likelihood that they are, indeed, relevant to epilepsy (i.e., related to an epileptogenic insult (30)). Interictal events and seizures projected into PC space were clustered using a Gaussian Mixture Model. We used the probability P(KA) that an event cluster was KA-specific to calculate an Hourly Epilepsy Index as a weighted average of the number of events with a P(KA) > 0.5. This value successfully estimates the probability that a subject had received an epileptogenic insult, in this case a KA injection. This algorithm can, in principle, be generally applied to chronic EEG recordings from either experimental animals or human patients, if a control data set is available for comparison. It can also provide rapid and quantitative scoring of interictal events to predict the likelihood that a subject is at risk for developing epilepsy even in the absence of observing electrographic or behavioral seizures. Importantly, the application of this algorithm to EEG recordings from animal models of epilepsy with low seizure burden (e.g. repeated low-dose kainate) may result in a reduction in the number of animals sacrificed in future studies.

### High-amplitude event detection is an inclusive starting point

Our approach to event detection using a two-threshold algorithm selected events of high amplitude compared to background signal, but did not, at the outset, impose limitations on event duration or morphology. This method is therefore intentionally inclusive of spike and other high-amplitude, non-spike-type events that may be physiologically relevant to epilepsy. Upon visual inspection of the signal and in comparison to a validated seizure- and spike-detection algorithm, we found that this method preferentially detects spike-type and seizure-like activity in the EEG signal (Figs 1 and 2), as well as other event morphologies, including artifact (Fig.3, cluster 8). That our detection method primarily selects for high-amplitude events does not negate the possibility that low-amplitude interictal events are relevant to epilepsy and epileptogenesis. Detection and analysis of low amplitude epileptiform events is an area for further inquiry (15).

### Selecting the appropriate number of clusters is flexible and application-dependent

We built on prior clustering approaches for EEG by considering how spike sorting may elucidate spike morphologies relevant to epilepsy and to differentiate these morphologies from non-epileptiform spike-like events observed in normal rodent brains (2,4). The high-amplitude events identified in records from SA- and KA-treated animals were projected into PCA space and then clustered using a GMM with expectation maximization. Similar to spike sorting in extracellular recordings (29), we found that the GMM provided a strong fit for the clustering of high-amplitude events in our dataset.

Nine clusters provided adequate separation between SA and KA-treated animals, though our analyses also found that this is, indeed, a fuzzy problem with many different options for cluster number selection depending on the features to be extracted. Spike morphology in extracellular recording is reflective of the geometry and physiological properties of individual neurons. Spike morphology in EEG may be reflective of a number of different parameters, including the underlying pathology of the spike, the spike location, the recording montage/referencing, individual patient and recording characteristics, and other features. Defining the number of clusters therefore requires careful consideration to prevent over- or under-estimation of the true number of spike morphologies.

We found that Cluster Belongingness and p-values for the separation between KA vs non-KA event morphologies improve as the number of clusters increase. However, very higher cluster numbers may be partitioning spike morphologies (clusters) based on individual animal or recording characteristics, rather than differentiating between KA and non-KA events. Selecting a smaller number (e.g. three rather than nine) of clusters for analysis does not provide statistically significant separation between KA and SA animals, so it is important to consider the tradeoffs in increasing and decreasing the cluster number for analysis (Figure 4). Through modeling using many different cluster numbers (from two to 25), we find that there is no clear “elbow” in the Negative Loglikelihood or Cluster Belongingness scores across our simulations. This indicates that there is not a hard and fast cluster number for this analysis. Still, we found that the p-value and effect size for the epilepsy index, and the Negative Loglikelihood values seemingly approached an asymptote by nine clusters (i.e. no major improvements at higher cluster numbers). Thus, we selected nine clusters for our analysis because it effectively separated KA from SA events while minimizing animal-specific effects.

We further explored event morphologies of individual clusters by specifically focusing on event amplitude and duration (Fig.5), as instructed by the descriptive spike definition of Chatrian *et al.* (1974). Events with a P(KA) > 0.5 (clusters 1-4, more likely to be found in KA than SA animals) exist within a space consistent with general definitions of spikes, that is events with high amplitude and short duration (the far lower left portion of the graphs in Fig.5A). As the P(KA) drops below 0.5 (clusters 5 through 9), the events begin to shift toward longer, lower-amplitude characteristics. It must be noted, however, that there is not a sharp demarcation in the amplitude and duration characteristics of spikes above and below the cutoff of P(KA) = 0.5.

In applying this analysis to other models of epilepsy, we see two major approaches to selecting an appropriate number of clusters, dependent upon the end goal of the analysis. One approach, similar to the approach that we have taken, is to identify the minimum number of clusters that provide significant separation between experimental conditions or a minimum cluster number that yield clusters of visually similar morphologies. A benefit to this approach would include the differentiation of noise and artifact from real epileptiform activity, as we see in cluster eight (Fig 3). Quantification of events could thus proceed relatively noise-free. This is similar to but more specific than an approach taken by (26), where spike morphologies in clusters that comprise less than 5% of the total spike count are removed from analysis. The considerations for a fine-scaled epilepsy index are different than this first approach, and would instead select a larger number of clusters for a higher-resolution model of epilepsy. Care must be taken, however, not to parcellate the data too far, resulting in over-fitting of the model.

### The Hourly Epilepsy Index: what it is, what it is not

We have introduced an Hourly Epilepsy Index as a summary descriptive variable of the likelihood of epilepsy in an individual animal, calculated as an average hourly event rate weighted based on the probability that an event morphology is found in KA- but not SA-treated animals. Given that kainate injection is but one model of epilepsy, we cannot yet claim that this is a true model that is applicable to all epilepsy models or etiologies. However, this index does begin to address some of the challenges of rodent models of epilepsy, namely that not all mice that receive KA-treatment will go on to develop epilepsy (30) and low seizure frequency may make it difficult to identify all epileptic mice, by providing relevant probability-based estimates of likelihood of an animal being epileptic.

There are several features of the Hourly Epilepsy Index that indicate that this may be a valuable model by which to analyze EEG data in epilepsy studies. At its simplest level, the Hourly Epilepsy Index provides a more robust separation between KA and SA animals than does raw event count (Fig 1C vs Fig 3D) by excluding events that are not likely to be epileptiform (events where P(KA) < 0.5) and weighting the remaining events (where P(KA) > 0.5) according to the probability that they are, indeed, epileptiform. The relevance of this new metric is further supported by data that reveals that the Hourly Epilepsy Index is significantly higher for the few KA animals with electrographic seizures than the average epilepsy index of SA-treated animals (Fig 3D). In further confirmation of this, we did not observe electrographic seizure activity in SA animals or KA-treated animals that fall within the 95% CI range of the Hourly Epilepsy Index for SA animals. However, these data do not fully confirm that all animals displaying a high Hourly Epilepsy Index have electrographic or behavioral seizures; further studies are required to confirm epilepsy.

Cluster one (Fig 3c, 5a) is of particular interest in considering the power of spike sorting by clustering for epilepsy analysis. These events are relatively infrequent across records, but are highly correlated with KA treatment (P(KA) = 0.99) and are very rare in SA animals. Visual inspection of these events confirms that they are, indeed, spike-like, fitting into a more classical definition of an interictal spike (18). Because these event types are almost exclusively seen in records from KA-treated animals, they are weighted more strongly in the epilepsy index than are events from cluster four, which show features of spiking (high amplitude, fast duration) but may also cause an expert scorer to pause on their definitive identity as an interictal spike.

The Hourly Epilepsy Index is not without caveats, however. Kainate is a widely-used model of TLE and epileptogenesis (30), but it is only one of many models. Other models, including pilocarpine, traumatic brain injury (TBI), genetic or viral models of epilepsy, and kindling, rely upon different molecular mechanisms of seizure induction and may therefore produce interictal events of different morphologies. Ultimately, this approach may be used to produce a library of spike templates across epilepsy models to further expand and refine the working definition of an interictal spike.

## Conclusion

Clustering of interictal spike provides insight into event morphologies relevant to epilepsy and epileptogenesis. Using this clustering method, we isolated epilepsy-like event morphologies that were then quantified to determine the probability that a particular animal is at risk for epilepsy or not, a variable which we have called the Hourly Epilepsy Index. This probabilistic classification method for interictal event waveforms provides a biomarker for risk and development of epilepsy, even in the absence of observing electrographic or behavioral seizures. Finally, by distinguishing spike morphologies that are preferentially present in the epileptic condition, we contribute to an unbiased understanding of the definition of interictal spiking and interictal spike morphologies, as compared with spike-like events present in the nonepileptic brain.

## Acknowledgements

We would like to thank Antoine Madar, Rebecca Willet, and Jun Zhu for helpful conversations during algorithm development.

